# EggVio: a user friendly and versatile pipeline for assembly and functional annotation of shallow depth sequenced samples

**DOI:** 10.1101/2022.04.23.489251

**Authors:** Benoit Marc Bergk Pinto, Timothy M Vogel, Catherine Larose

## Abstract

We introduce a homemade pipeline allowing to improve the quality of the metagenomic annotations carried out when using shallow depth metagenomic datasets. The main motivation being to be able to quantify more precisely, with greater certainty, the genes involved in bacterial interactions. The limitation in our experimental design is that we use a sequencing technique with a low throughput (miSeq) compared to the metagenomic standard (hiSeq) because we carry out a fairly large sampling (almost a hundred samples) in time series. This methodological constraint from our study means that the assembly of the sequences is not very exhaustive (less than 50% of the sequences manage to be assembled). In this chapter, we will therefore present a new pipeline designed to specifically deal with such kind of data. We used co-assembly and a sequence annotation strategy in order to recover the sequences that could not be mapped on the assembled contigs. In addition, in order to avoid adding too much noise, when rescuing reads, we have built an algorithm to define a threshold of e-value based on the noise of the sequence annotation learned from sequences mapped in the assembly.

We have selected several recent tools known to be effective for assembling, mapping and annotating these data. In addition, this pipeline was also built in order to be very user-friendly in terms of installation. In this idea of reproducibility, accessibility and transparency, we have designed an installation script to allow each user to install each tool required for the pipeline in a simple and reproducible way. Regarding the performances of this pipeline, we were able to show that the expected error rate (False discovery rate) for the annotation was close to 5%. Finally, we also used an actual dataset from a bioremediation site and showed that the representability of the samples seemed much better when we used our pipeline than when we used a classic metagenome assembly strategy.

## 2 Introduction

Metagenomic approaches are useful for investigating both the diversity and functioning of environmental microbial communities. Over the last decade, several tools and workflows have been released to assemble and analyse these datasets (*e*.*g*. Li et al. 2016; Bankevich *et al*. 2012; Wood and Salzberg 2014; Buchfink, Xie, and Huson 2015; Menzel, Ng, and Krogh 2016). Metagenomic data (derived from shotgun sequencing of total extracted DNA) has recently been used to assemble putative genomes, called metagenomic assembled genomes (MAGs), to determine taxonomy and to identify metabolic functions of microbial communities. Sequence reads are assembled into contigs, which improves the accuracy of annotation, and binned into MAGs. Both the contigs and the bins (putative genomes - MAGs) can provide more accurate taxonomical annotations. The accuracy of the processes suggested for assembling the reads in contigs and for binning them are dependent on sequencing depth; if it is too shallow, the coverage of the contigs will be low and only a small fraction of reads will be recruited into the assembly. As a consequence, the results might be less reliable and other strategies must be used.

There are several methods for improving the assembly into contigs and for the subsequent binning into MAGs. Co-assembly is one strategy that has been developed to improve the assembly quality when sequencing depth is limited. If the metagenomic dataset is composed of several samples that are known to contain similar bacterial communities, they can be pooled during the assembly step to increase the sequencing depth and improve the quality of the assembly. In this case, reads from different samples can be used to create a contig. The subsequent binning of contigs has often in the past relied on nucleotide frequency discrimination, although the binning could be improved by using differential coverage of the reads in the contigs in the different samples. Co-assembly of the contigs from multiple samples could, therefore, improve the quality of MAGs retrieved by increasing the discrimination among taxonomically related species or strains. For example, Delmont and Eren (2018) retrieved more MAGs from the Tara ocean dataset, including several strains of *Prochlorococcus*, than found in the original publication (Sunagawa et al. 2015) by using both co-assembly of the contigs and differential coverage for the binning. Co-assembly can also be used to track changes in taxonomy and functions in time-series datasets. If the genes of interest are abundant, shallower sequencing technologies other than hiseq (*e*.*g*. miSeq) can be used and co-assembly can increase the assembly quality. However, pipelines to deal with these specificities (time series, shallow depth sequencing, low assembly) are currently missing.

In this paper, we introduce a new workflow (“Eggvio”) designed to annotate and analyse metagenomic datasets of any size (but optimized for shallow sequencing datasets). Co-assembly was coupled to a read annotation strategy to rescue reads that were not easily mapped back to the assembled contigs. In addition, an algorithm was built to define an e-value threshold based on the noise of read annotation derived from the reads used in the assembly. Several recent tools to carry out the assembly, mapping, binning and annotation were selected to build a user-friendly pipeline. Pipelines for metagenomics generally involve the installation of many tools or dependencies and require some previous informatics knowledge. To improve the reproducibility, accessibility and transparency, we designed a script that allows every user to install all the tools needed for the pipeline.

## 3 Material and methods

EggVio is a flexible pipeline optimized for co-assembly, binning and annotation of low depth MiSeq samples (<10^6^ reads per sample). Due to the computational needs for read annotation, it was designed to be used on a linux server with a batch slurm queue for job submission. It is coded mainly into bash scripts to process fastq data starting from quality filtering to functional annotations. R (R Development Core Team 2011) programming is used for merging annotations from contigs and reads that did not map back onto the assembly and also to determine the e-value threshold. The programs used to carry out every step of the pipeline are shortly summarized in the flowchart shown in Figure 1. In addition, an installation script (EggVio_install_tools.sh) is provided for most of the tools that do not require any root privileges for installation (this excludes the assembler, megahit). The user needs to download the annotation database for eggnog mapper and the taxonomic database for contig taxonomic annotation by kaiju separately. The databases were not included in the installation, because multiple annotation databases are available and the user can choose which one to download themselves. As a guide, we included both steps in the wiki pipeline (https://gitlab.com/R_addict/eggvio/wikis/Download-and-installation-of-EggVio). The scripts for the pipeline are freely available on GitLab (https://gitlab.com/R_addict/eggvio) under the MIT open source license.

**Figure 1:**
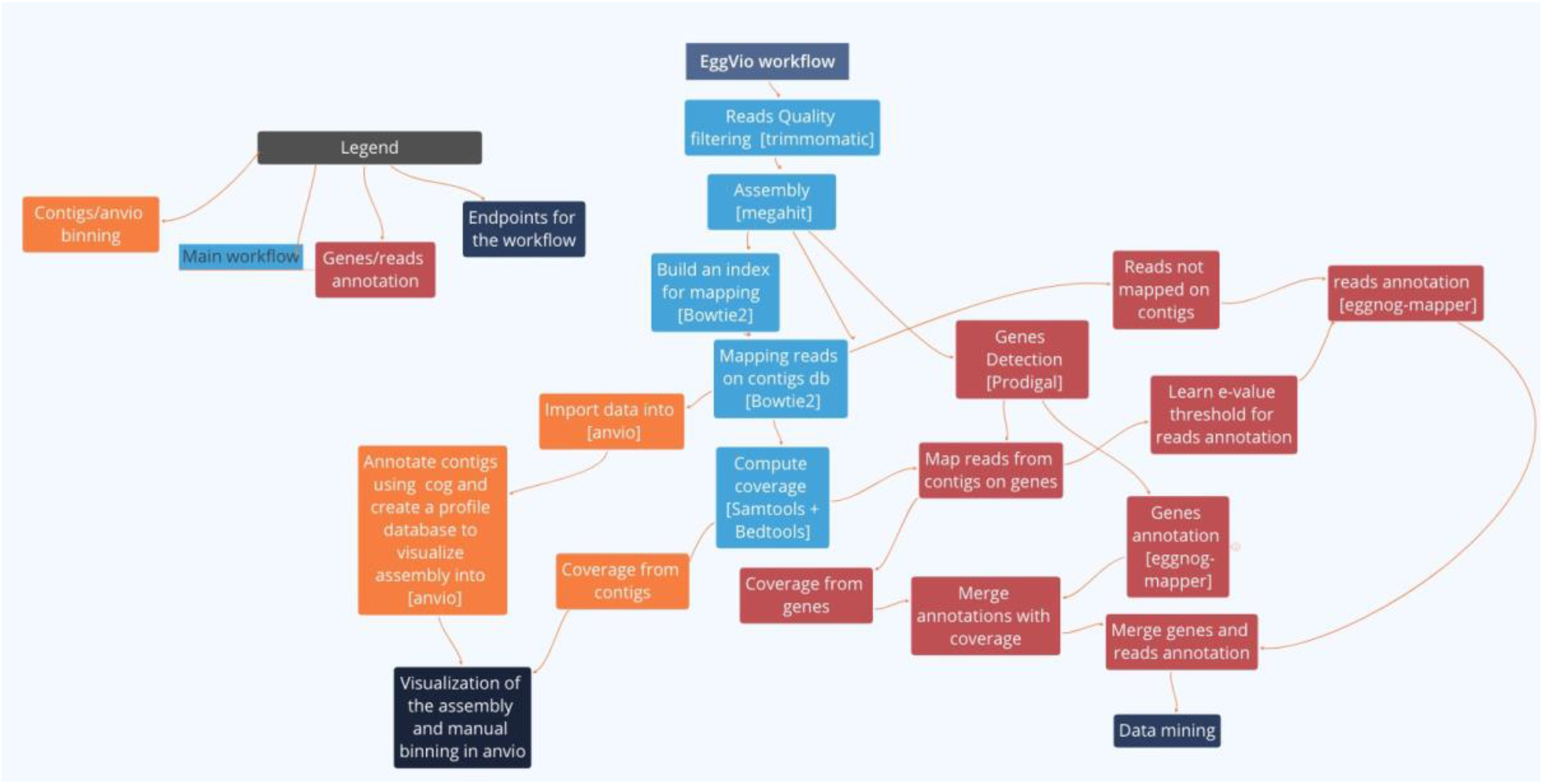
Summary of the workflow. The workflow is divided in three subparts illustrated by different colors. The main workflow (light blue) carries out steps which are mandatory to both MAG binning with anvio (orange) and functional annotation (red) using eggnog-mapper. Tools used to carry out every step of the pipeline are written between brackets inside the box. If no tool is given, then it means that the whole step is carried out using custom R or bash scripts without the need of any other tool.

### 3.1 Description of the steps and citation of the tools used in the pipeline

The Eggvio pipeline can be applied to raw metagenomic illumina reads for MAG assembly and refinement in anvio or for functional annotation using a hybrid method (assembly and read annotation). The main steps of the pipeline are represented in Figure 1 in light blue. The first step is quality filtering using trimmomatic (Bolger, Lohse, and Usadel 2014) followed by assembly using megahit (D. Li et al. 2016). The assembled contigs are then renamed using anvio (Eren et al. 2015) and filtered by length (optional) to discard contigs that are too small for binning and gene prediction steps. Read mapping onto the assembly is computed using Bowtie2 (Langdon 2015). If the user is interested in assembling MAGs, the next steps of the pipeline consist in taxonomic annotation of the contigs using kaiju (Menzel, Ng, and Krogh 2016). These data can then be imported into anvio for binning of MAGs and assembly visualization. For functional annotation of the metagenomes, several other steps are needed and are described in the next section with a focus on the algorithm used to learn the e-value threshold for read annotation.

### 3.2 Description of the learning algorithm to estimate the e-value threshold for read annotation

After co-assembly of the different samples of the dataset (Figure 2 A), gene detection on contigs (Figure 2 B) is carried out using prodigal (Hyatt et al. 2010). The coverage is then computed using Bowtie2 (Figure 2 C) and results are converted into counts using a custom bash script relying on functions from samtools (H. Li et al. 2009) and bedtools (Quinlan and Hall 2010). The genes and the reads which mapped onto them are then functionally annotated (Figure 2 D) by eggnog-mapper (emapper version: emapper-1.0.3-3-g3e22728 emapper) (Huerta-Cepas et al. 2017) using the diamond (Buchfink, Xie, and Huson 2015) mode with the eggnog orthology database (DB version: 4.5.1). These gene annotations and their respective mapped reads are compared to identify false positive (FP) annotations. If the read annotations differ from the genes they are mapped onto (considered as the ‘gold standard’ annotation), this is considered as a false positive (Figure 2 E). Based on these data, the e-value threshold (E.T.) learning algorithm is used to define a suitable threshold for read annotation where the percentage of expected FP in the annotations considered as significant would be p-value <= 0.05.

**Figure 2:**
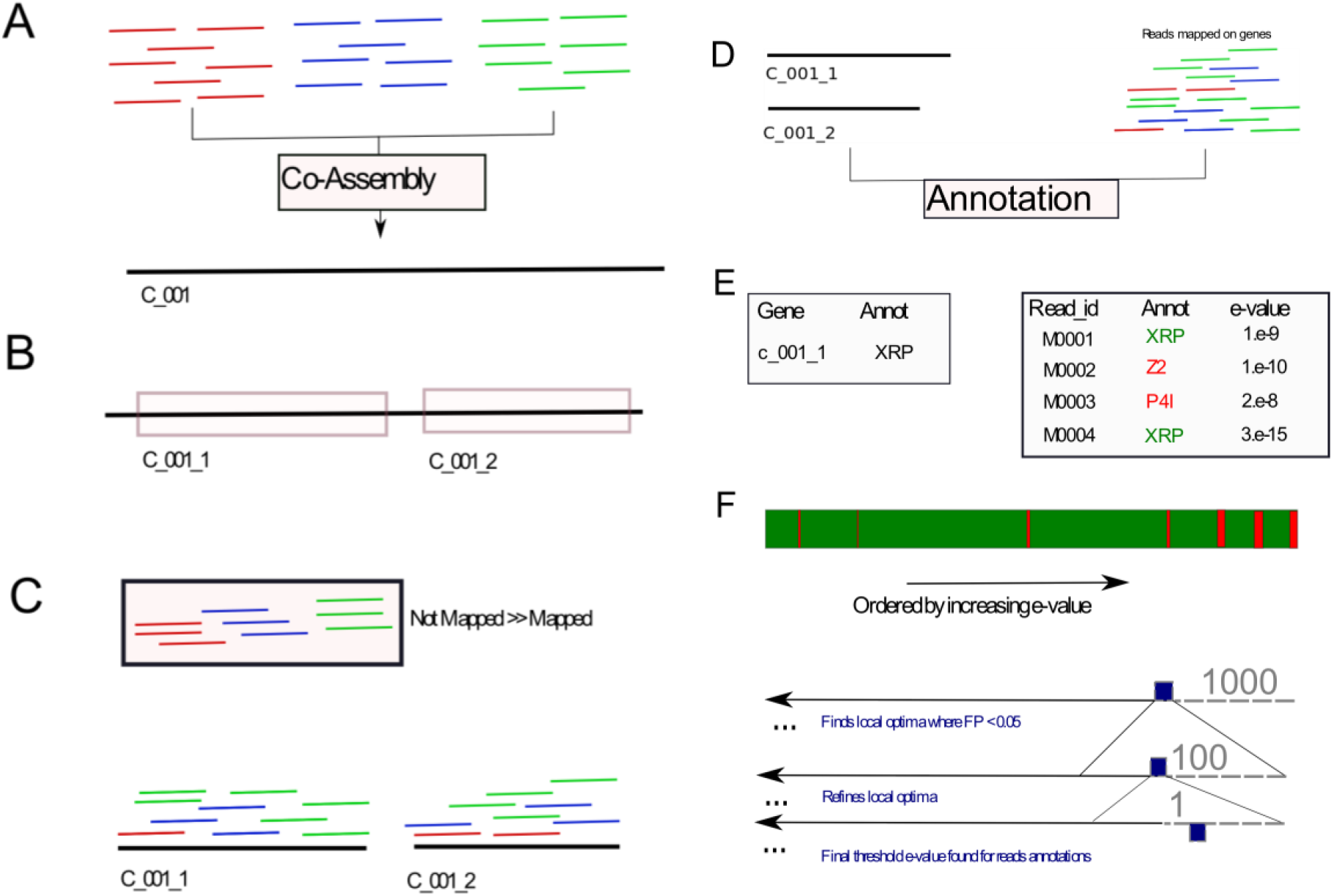
Summary of the different steps of the EggVio pipeline needed for read annotation. **A**. The reads from different samples from the dataset are co-assembled into contigs **B**. Genes are detected on the contigs and extracted in fasta. **C**. Reads mapped on the contigs (previous section) are mapped on the genes and their respective coverage across the dataset is computed. **D**. Reads which mapped successfully on the genes and the genes themselves are functionally annotated. **E**. The annotations of the genes (considered as a ‘gold standard’) are compared to the annotations of their respective mapped reads. If the annotations of reads and genes are different, the read annotation is considered as spurious (False Positive = FP) and if they are identical, then they are considered as true positives (TP) **F**. The algorithm to learn the threshold is a greedy algorithm based on successive refining steps. It works on a (decreasingly) ordered vector of e-values from read annotation with a corresponding vector returning the information whether the corresponding e-value returned a TP or a FP annotation. The algorithm will compute the percentage of FP starting from the whole dataset and then by removing iteratively the highest 1000 e-value annotations. When it finds the local optima, it will start computing the same statistics, but starting from the interval identified as optimal and refine it by removing iteratively 100 annotations and then for the last refinement step only 1. This e-value returned will then be used to annotate the reads that did not map onto genes with an expected probability of FP<0.05.

This algorithm is written in R and is presented below.

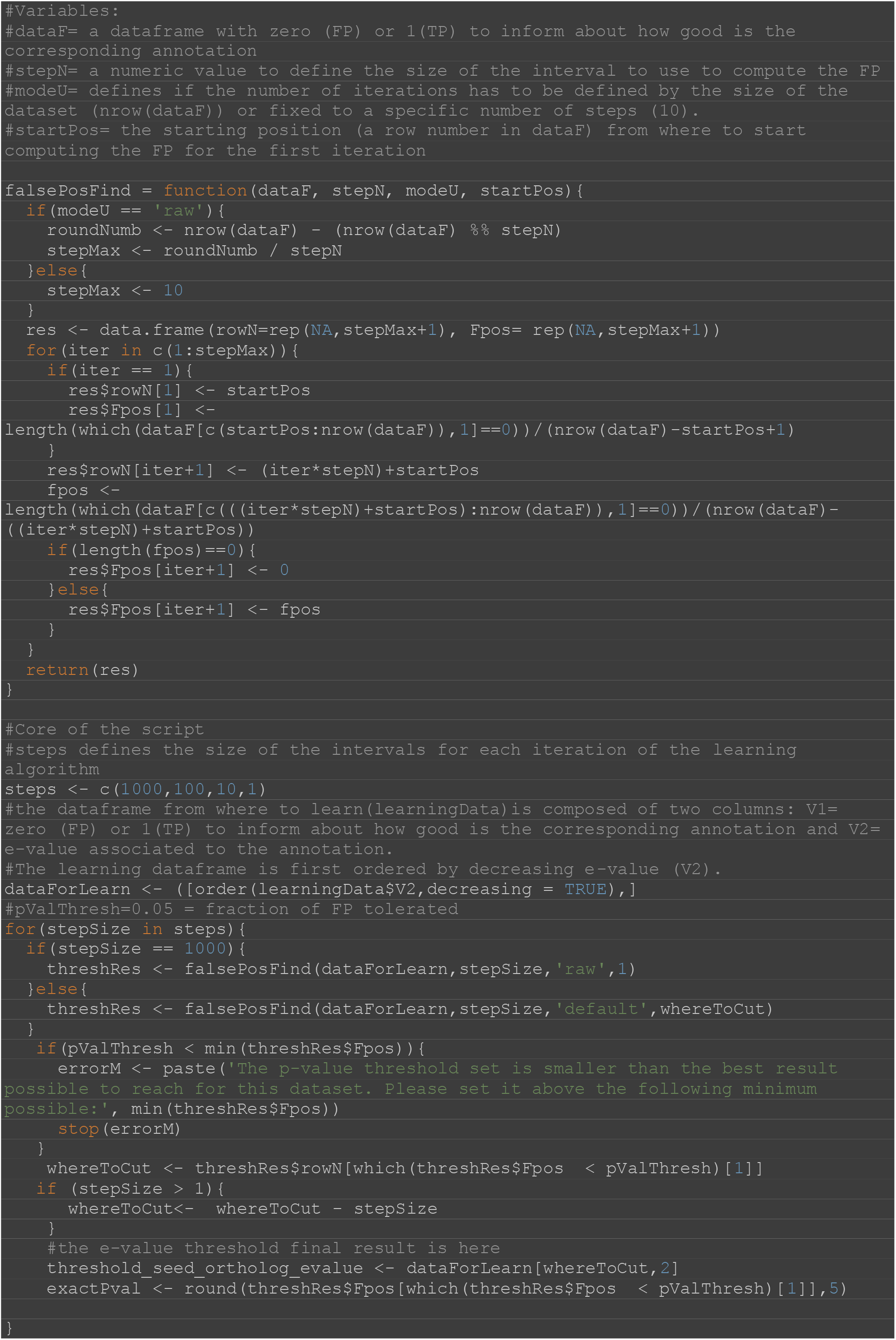

This function is designed to find an e-value threshold such that after filtering, the false discovery rate 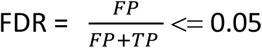 (where FP = False Positive and TP = True Positive)

At the same time, we would like to minimize the rejection of correct annotations. To meet both criteria, the function orders the dataset by decreasing e-value (the least significant e-value has the row name 1 in the ordered dataset). Then, the script calls the function “FalsePosFind” to compute the FDR with a threshold set every 1000 annotations. Since the data are ordered, the first threshold that meets the criteria FDR < 0.05 will be the best solution, as it will preserve the highest amount of TP. Once this position is found, the loop will iterate the function “FalsePosFind” a second time, but starting from the position located 1000 annotations higher than that initially found. The optimization serves to minimize the amount of TP rejected as false negative (FN) and to refine the threshold such that it is less restrictive by computing FDR every 100 annotations starting 1000 annotations away from the local optima found on the previous iteration. In total, 1000/100 = 10 FDR will be computed and then the best optima (first optima found) will be selected and refined further with intervals of 10 annotations and finally 1 annotation. This e-value threshold will then be used to annotate the reads not mapped on the contigs using eggnog-mapper.

### 3.3 Benchmarking the EggVio pipeline

Presentation of the dataset used in the study:

To test how our pipeline could enhance the annotation of shallow sequenced datasets, we used one of our in-house datasets from a polluted site bioremediation project (MISS). It consisted of a time series of 30 samples tracking chlorinated compounds in a polluted ground water site. After the injection of organic carbon to induce the biodegradation of chlorinated solvents in the groundwater (C1), three other samples were taken one month apart (C2, C3 and C4). Six replicates were collected for each time point. The biodegradation of chlorinated compounds was evaluated at each time point and qPCR analysis was carried out in order to evaluate the abundance of the pceA gene coding for an enzyme involved in reductive dechlorination of tetrachloroethene (PCE) to trichloroethene (TCE).

### 3.4 Evaluation of the threshold learning algorithm

We first used this dataset to evaluate how our threshold learning for read annotation based on assembled data would perform. We computed the threshold for every annotation (gene id, gene name, Kegg orthologs = KO and Gene Onthology = GO) to compare the thresholds returned. We then focused on the KO annotation to test how learning on a subset of the dataset (since we could not assemble the whole dataset) would affect the threshold and FDR estimate. To test this, we randomly subsampled the learning dataset to predict the threshold and then observed how the FRD was affected when computed on the whole dataset.

### 3.5 Comparison of the sequencing results and the qPCR results on the tracking of the gene pceA

We also used this dataset to compare how the hybrid annotation (genes and reads annotations) would affect the results as compared to gene only annotation using the intermediate results from EggVio. Both annotations (hybrid and genes only) were then normalized by the RPKM method using the R package GenomEnvironR (github: https://gitlab.com/R_addict/genomenvironr). The percentage of sample being annotated as genes and as reads was then investigated for every sample. A NMDS of the samples was carried out for both types of annotation at the KO (Kegg orthologs) level. These sample representations were compared to an NMDS plot of the samples that included chemical data of several chlorinated compound concentrations measured in the water. Given that measurements were missing or below detection level for some of the chlorinated compounds, we only used PCE, TCE and cis-DCE in the analysis.

Finally, we determined whether read annotation using the hybrid EggVio approach would improve classic methods by comparing the abundances of pceA genes observed in the contig assembled data versus the hybrid annotation and the qPCR data. To determine whether this approach could improve the overall significance between biological and chemical data, we correlated gene abundances with environmental variables related to pceA activity (PCE and TCE).

## 4 Results

### 4.1 Benchmarking the EggVio pipeline

#### 4.1.1 Summary of the assembly and reads annotation

In total, the assembly recruited only 11% of the reads (915788 reads from 8001127 reads in total) used in the co-assembly and generated over 319458 contigs. The N50 of the assembly was 593 bp of contig length. During the mapping, 100% of the assembly could recruit at least one read of coverage showing that no artefact contig had been generated during this step. In addition, most of the assembly displayed a coverage of 2 minimum (99% of bins of size 10 bp displayed a coverage of 2 or more).

The number of reads annotated by their annotation in their respective contig assembly was high for some samples (up to 34%), but heterogeneous, with some samples being annotated below 1% (Figure 3). Individual read annotation provided more homogeneous results, with all the samples having more than 10% and up to 26% of their reads annotated (Figure 3). For the hybrid method, the lowest percentage of annotated reads was around 15% and reached up to 65%. On the other hand, the range of possible annotations decreased dramatically when contigs were used. In other words, the annotation variability at the seed eggnog ortholog level (composed of a unique gene and its respective genome taxonomy id) increased significantly for the reads compared to the contigs. The gene annotation returned 18613 possible genes from the contig annotation compared to 408618 different genes (∼ 22 times more) for the read annotation.

**Figure 3:**
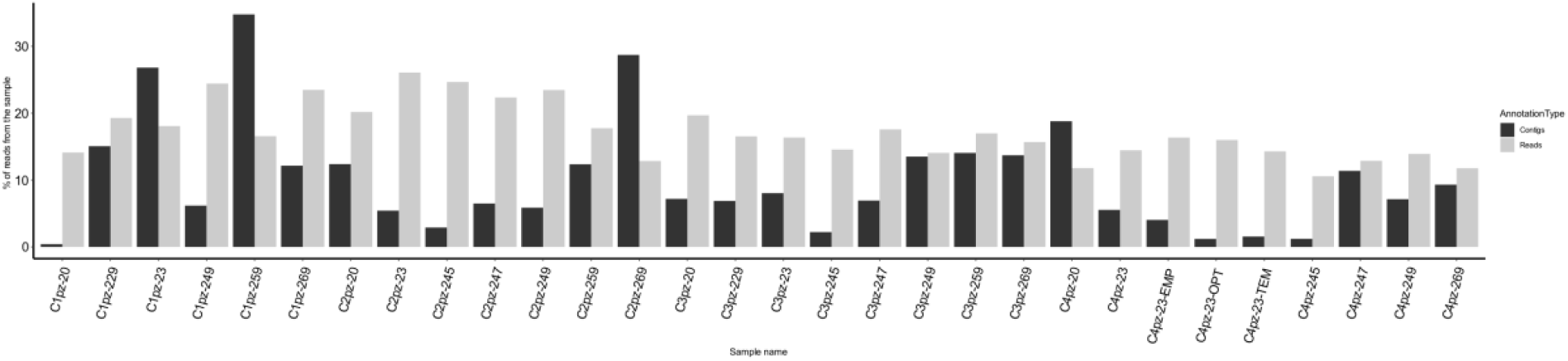
Summary of the percentage of reads annotated successfully (seed eggnog ortholog level) with the hybrid method from EggVio. The annotations derived from the genes predicted on contigs are represented as black bars and the annotations derived from direct read annotation are represented as grey bars.

#### 4.1.2 Evaluation of the read annotation threshold learning algorithm

In order to evaluate how our learning threshold performed, we first ran the algorithm on the fully assembled data. Then, we assessed how random subsampling of the training dataset would impact our learning threshold and its ability to keep the amount of false positive below 0.05. To do this, we subsampled the mapped reads to smaller fractions (respectively 80%, 50 % and 10% of the original dataset) 1000 times and ran the threshold analysis. The threshold results were then compared to the amount of True Positive (TP) and False positive (FP) in the original dataset at different annotation levels (gene ID, gene name, KO and GO, Table 1). An FDR = 0.05 was obtained for all the annotation levels except for Gene ID, where the error was too high with a threshold at FDR = 0.34. By default, the algorithm computes an e-value threshold where the False discovery rate (FDR) is below 5% of False positive (FP), i.e. FDR = 0.05. For each threshold, the number of true positives (TP) is also given. The fraction of correct annotations rejected with this threshold is also given (fraction of correct rejected) as well as the fraction of false annotations successfully rejected (fraction of false rejected). These two fractions are used to determine the sensitivity of the annotation.

**Table 1:** Results of the e-value threshold learning algorithm for the MISS dataset. False discovery rate (**FDR)**, false positive (**FP**), true positives **TP**, false negatives **FN**, and true negatives (**TN**)

| ANNOTATION | FDR | THRESHOLD<br>E-VALUE | FN | TN | FP | TP | FRACTION OF<br>CORRECT<br>REJECTED | FRACTION OF<br>FALSE<br>REJECTED |
| --- | --- | --- | --- | --- | --- | --- | --- | --- |
| GENE<br>ID | 0.34 | 1.8e-27 | 0.52084 | 0.47916 | 0.34001 | 0.65999 | 0.6348 | 0.75634 |
| GENE<br>NAME | 0.05 | 2e-08 | 0.82506 | 0.17494 | 0.04995 | 0.95005 | 0.02547 | 0.09535 |
| KEGG<br>ORTHOLOGY | 0.05 | 3.2e-15 | 0.88202 | 0.11798 | 0.04988 | 0.95012 | 0.26222 | 0.4752 |
| GENE<br>ONTOLOGY | 0.05 | 3.5e-17 | 0.83239 | 0.16761 | 0.04998 | 0.95002 | 0.30443 | 0.62618 |

Based on the annotation level, the threshold is variable for the same FDR (0.05). The highest threshold found for an FDR of 0.05 was for gene name annotation (2e-08) and the lowest was for GO terms annotation (Table 1). The vast majority of the annotations rejected are FN (above 80% of the rejected annotations during the learning) (Table 1). This feature is expected since the e-value distributions of the correct and incorrect annotations for KO (Figure 4) overlap. If we compare their medians (dashed lines on Figure 4), the e-value median of the correct annotations is much smaller (<1e-20) than that of the incorrect annotations (>1e-15). As a consequence, the percentage of correct annotations rejected is smaller (∼26%) than that of the incorrect annotations (∼47%) (Table 1).

**Figure 4:**
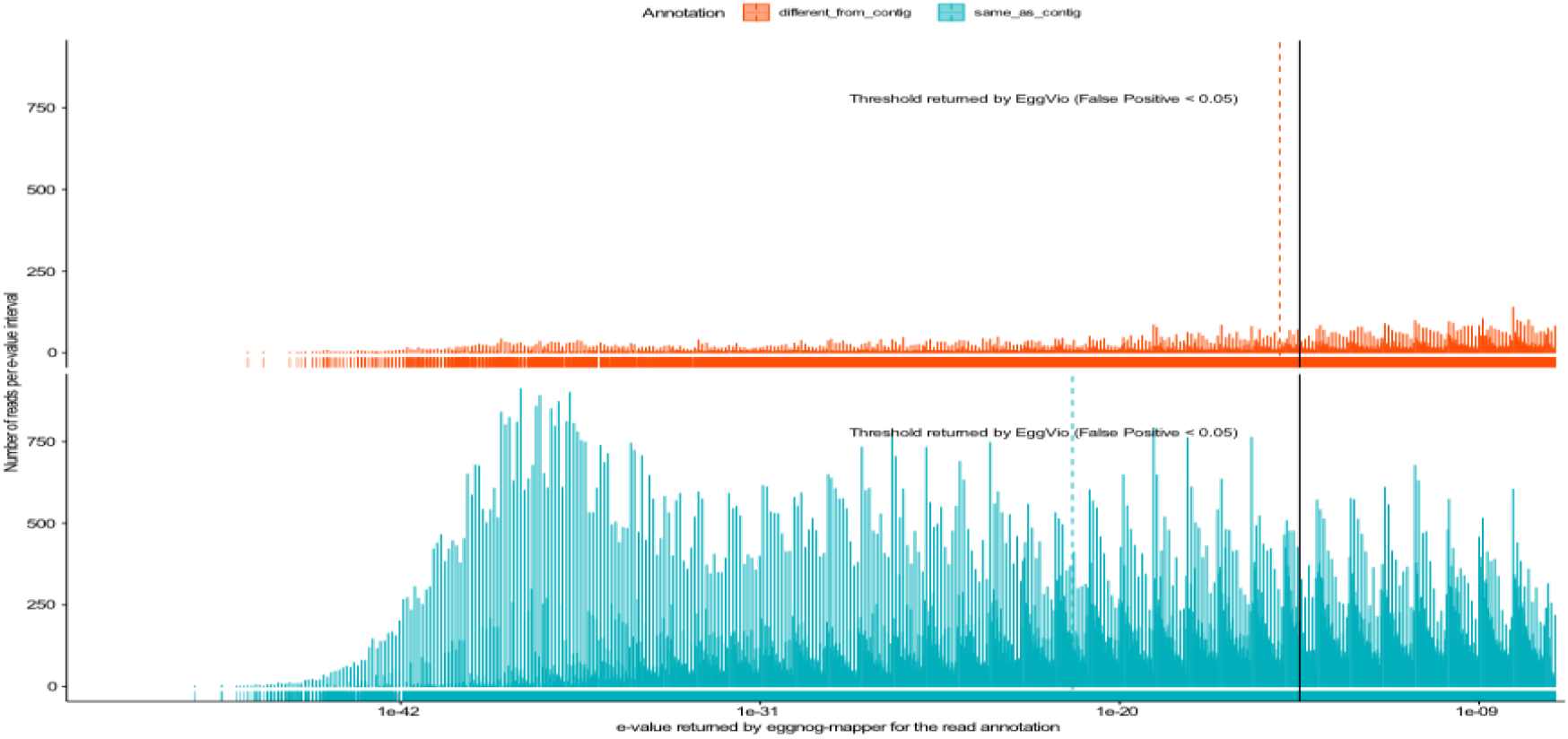
Histograms of the distribution of e-value for the reads annotations from the reads mapped on the genes predicted from the contigs of the assembly. The vast majority of the read annotations are identical as the ones from the genes (blue green) at the opposite of the spurious annotations (red) different from the genes. The vertical black line represents the threshold returned by the learning algorithm from EggVio where the amount of spurious annotation represents less than 5% from the total number of annotations accepted with this threshold. The dashed vertical lines represent the median e-value of their respective distributions.

We then evaluated the impact of dataset subsampling on the e-value threshold estimate for read annotation and our FDR estimate for the whole dataset. After plotting the results of 1000 subsampling at different percentages of the dataset, we observed that the e-value estimate became more variable at smaller subsampling sizes (Figure 5) and the boxplots tended to deviate from the true estimate toward smaller e-values as they were moved toward the bottom of the y axis (Figure 5).

**Figure 5:**
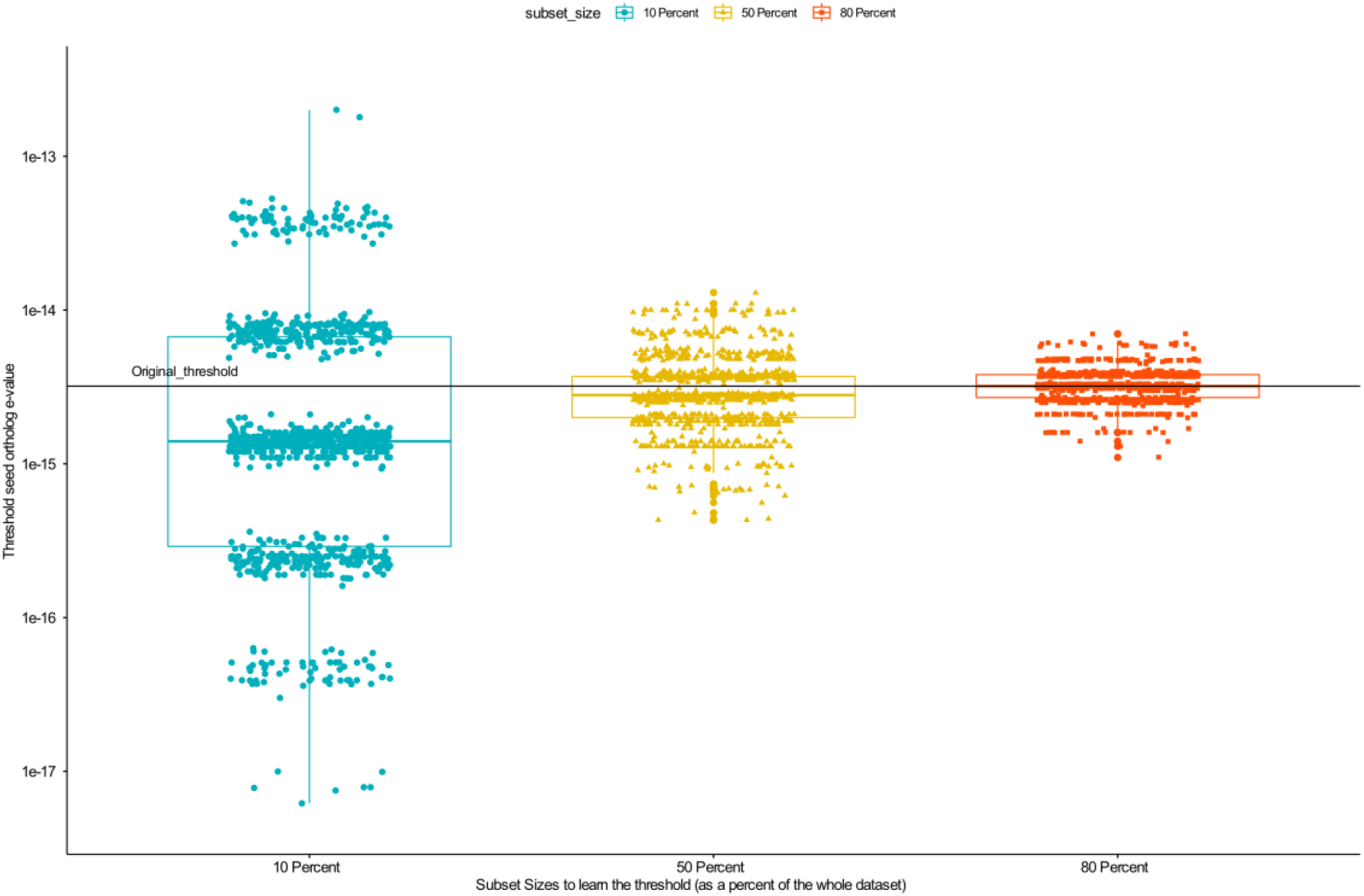
Effect of subsampling on the learning of the e-value for the read annotation. Each dot represents the threshold e-value returned by the algorithm based on its learning on a subset of the annotations (represented on the x axis as a percentage from the total dataset) from the reads mapped on the assembly genes.

This trend was confirmed by the amount of TP observed for each of the e-value thresholds returned for the different subsampling. The second quartile of the boxplots was generally above 0.95 (Figure 6) for the subsampling replicates, indicating that for more than 50% of the subsamples, the threshold returned was enriched with correct annotations and thus showed an FRD <0.05. In addition, the replicates with the lowest amount of TP were always above 0.945, and FRD <0.055 was observed in the worst case during the subsampling experiments (Figure 6).

**Figure 6:**
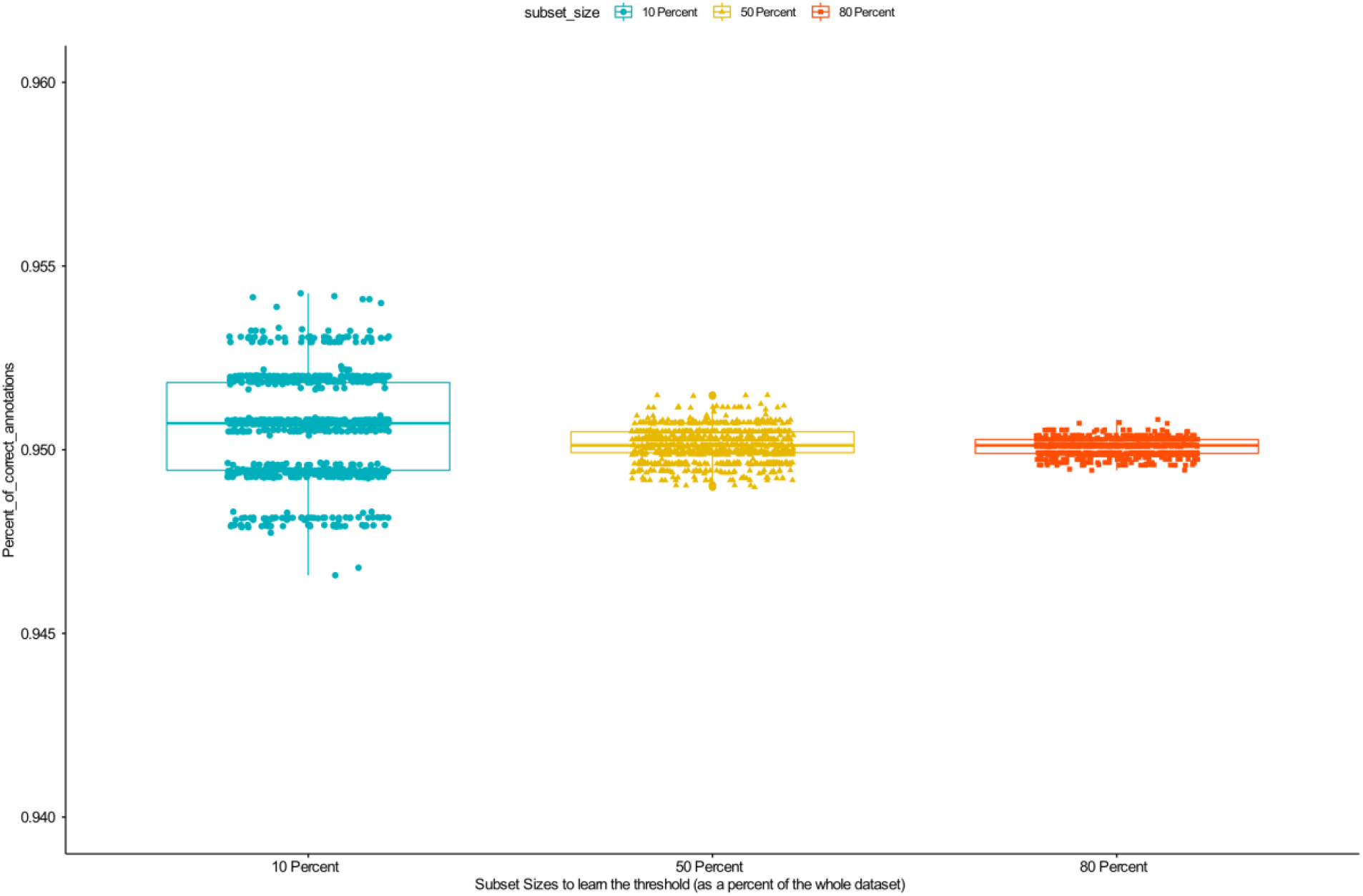
Effect of subsampling on the amount of true positive (TP) after filtering. Each dot represents the amount of TP detected in the annotation of the reads mapped on the genes (whole dataset) when the learning was performed on a subset of the annotations (represented on the x axis as a percentage from the total dataset) from the reads mapped on the assembly genes.

### 4.2 The effect of the read annotation on the dataset representability and the detection of pceA genes

We investigated how the NMDS representations of the MISS dataset based on the metagenomic data agreed with the NMDS representation based on the chemistry (Figure 7). The NMDS from both types of annotation (assembly and hybrid) clustered more together at a given sampling time (*e*.*g*., C1 with samples C1) than between samples from different sampling times, but their grouping at a finer scale showed differences. For example, the sample C1pz-20 was far from all the other samples from the dataset when considering only the assembly annotation (Figure 7 A) but did not display any chemical features on the chemistry NMDS that could explain its position (Figure 7 C). This sample had the lowest annotated coverage (<1%) in the assembly (Figure 3). We also observed that like in the NMDS of the chemistry, the samples from sampling time C4 were more densely clustered together on the NMDS from the hybrid annotation (Figure 7 B) than for the assembly only (Figure 7 A). A last trend to observe is that at the opposite of the chemistry NMDS representation, the samples C3 are totally apart from the samples C2 and separate C2 and C4 samples in the assembly NMDS (Figure 7 A). The NMDS from the hybrid annotation agreed more with the continuum between C2, C3 and C4 and led to C2 and C3 samples to be partially intermixed (Figure 7 B).

**Figure 7:**
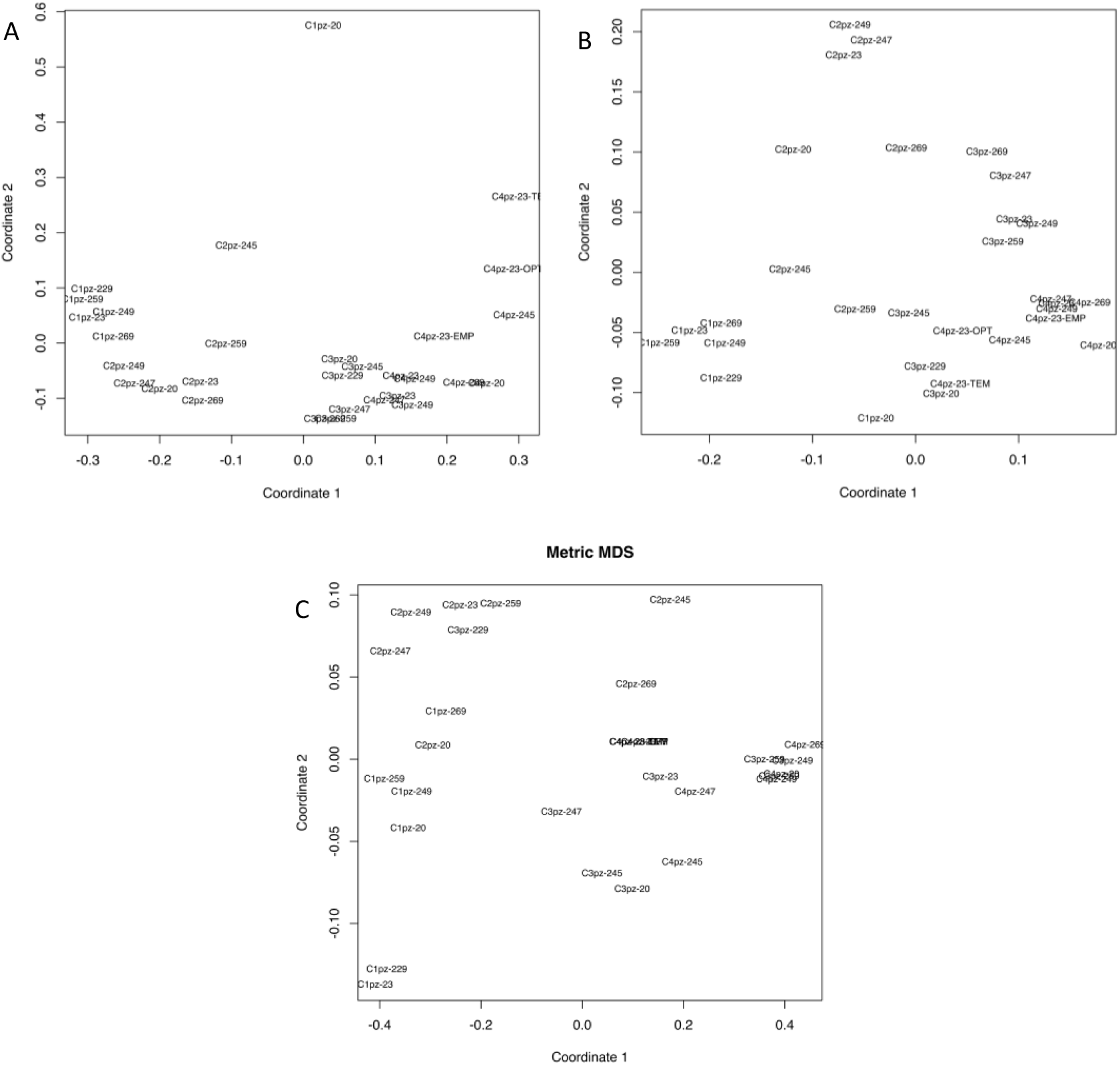
NMDS of the different samples from the MISS dataset based on their respective normalized annotations (Kegg orthologs) established on the annotation of: **A** = genes predicted from the contigs assembly only **B** = genes predicted from the contigs assembly and annotations of reads not mapped on the assembly. **C**. = NMDS based on the chlorinated compounds measured (PCE, TCE and cis-TCE).

The last result generated is the follow up of the relative abundance of the pceA genes. The abundance of these genes at different sampling times was determined for the different metagenomic annotations and quantified using qPCR (Figure 8 A, C and E). We observed that the sequencing data totally disagreed with the qPCR data (Figure 8 A versus C and E). The maximum amount of pceA quantified using qPCR was detected at time 1 and 4 at the opposite of the metagenomic data where the maximum observed quantities were detected at t2 and t3 (Figure 8 A versus C and E). Interestingly, we could observe that in the metagenomic data, the vast majority of the signal was coming from the assembly (>90%) by comparing the plots from the assembly to the hybrid annotation (Figure 8 C versus E). We could not detect any correlations between pceA gene quantification and PCE or TCE concentrations in the samples (data not shown) but we could detect a significant negative correlation between the quantification of pceA genes in metagenomic data and the ratio of the PCE over TCE concentration (Figure 8 D and F). interestingly, the detected correlation was more significant when computed using the hybrid data quantification (p-val=0.018) instead of the assembly annotations (p-val=0.046) alone (Figure 8 D and F).

**Figure 8:**
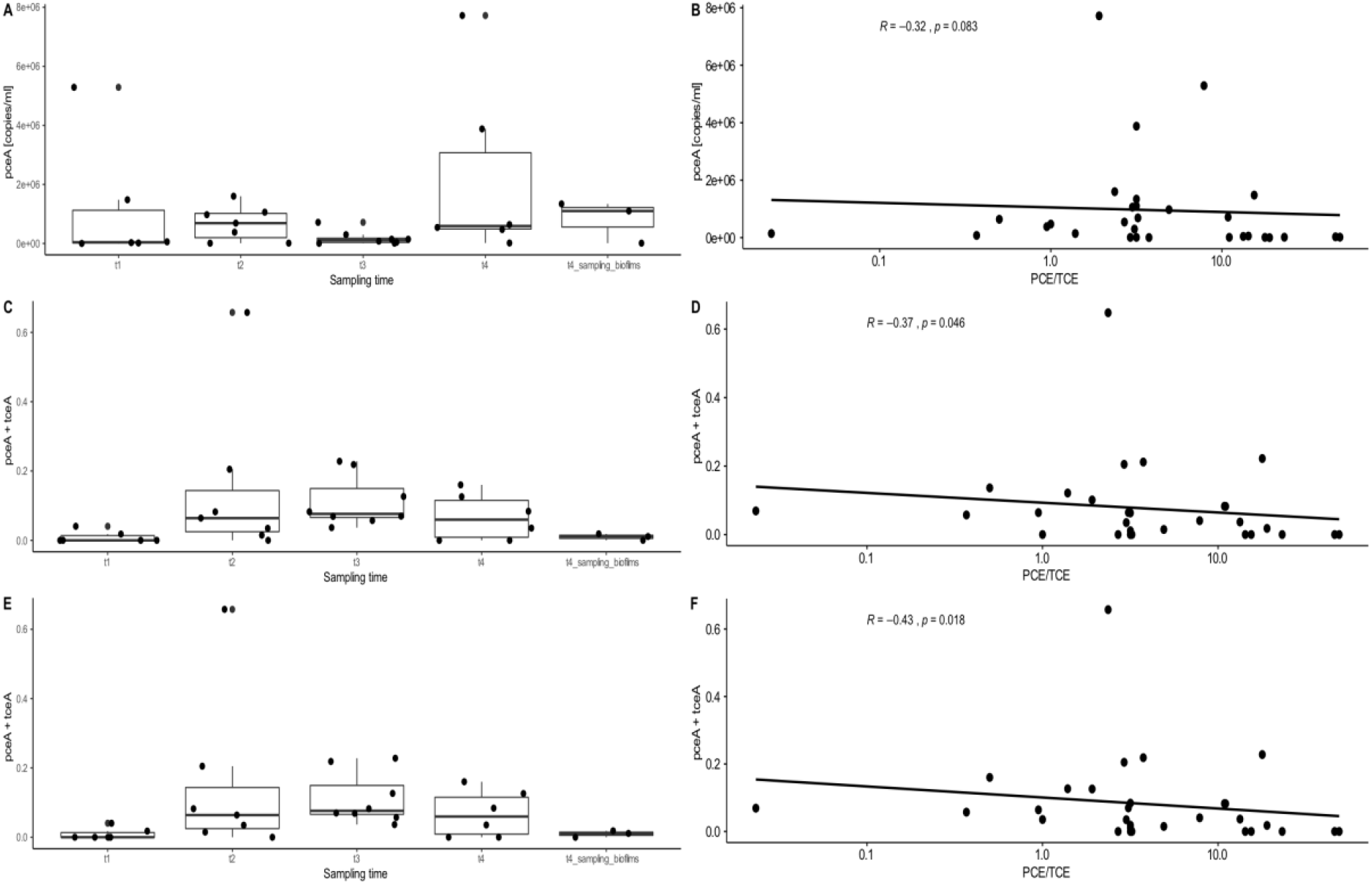
Comparison of the abundances of the genes pceA detected using qPCR (**A**), sequencing with gene retrieved from the assembly alone (**C**) or with the reads annotation added (**E**). These abundances detected with those methods have then been correlated to the ratio of the target of the gene (PCE) and its end-product (TCE) using the qPCR data (B), the sequencing with gene retrieved from the assembly alone (**D**) or with the reads annotation added (**F**).

## 5 Discussion

### 5.1 Assembly missed meaningful information in shallow depth datasets, but can be complemented by rescuing reads

The percentage of reads recruited in the assembly varied widely across samples (Figure 3), with an average of 11% of reads recruited and the amount of gene ids detected in the assembly were 22 times lower than in the read annotations. This is lower than what was observed in a study where full datasets were assembled and 10% to 30% more annotations were retrieved using an assembly free method (Anwar et al. 2019). The undersampling of the dataset was further supported by the NMDS representation that showed that sample C1pz-20 was the most unique sample based on the assembled data (Figure 7 A). However, this sample had the lowest coverage (<1%) in the assembly (Figure 3). After applying the hybrid method, the NMDS could be corrected (Figure 7 B) and showed a representation of the samples that was much more in accordance with the chemical dataset (Figure 7 C). In addition, the hybrid method increased the sensitivity of correlation detection between the relative abundance of pceA genes in the metagenomes and the concentrations of PCE over TCE (Figure 8 E versus F). This negative correlation was not detected in the qPCR data (p-val>0.05). We interpreted this correlation as PCE degradation occurring prior to sampling time rather than an instantaneous measure of the PCE degradation potential of the bacterial community (rather detected by metatranscritomics). Thus, at lower PCE/TCE ratios, a higher abundance of pceA genes could be interpreted as a selection for organisms able to degrade PCE into TCE in the microbial community.

### 5.2 EggVio E.T. algorithm can accurately quantify the noise added when rescuing reads for annotation

By randomly subsampling the learning dataset to evaluate how the e-value threshold estimate would affect the estimation of the FDR at the KO annotation level, we showed that in the worst case, the FDR would only be impacted by 5% leading to an accuracy (= TP/(FP+TP)) of 0.945 (Figure 6). We also observed that the recall (= TP/(TP+FN) = 1 - FRACTION OF CORRECT REJECTED) was above 69% if we exclude the gene id (seed eggnog ortholog). These estimated performances can be compared to other read annotation tools such as miFaser (Zhu et al. 2018), recently released and based on a custom high quality database using a custom score modeled after the HSSP metric for function transfer between full-length proteins (Schneider, de Daruvar, and Sander 1997). Based on reads generated from their database, they assessed that the accuracy of miFaser could reach 90% and the recall 50% (Zhu et al. 2018). However, since the databases are different and do not use the same criteria, comparisons are difficult to perform.

Concerning the lowest level of annotation accessible by eggnog-mapper (gene id = seed eggnog ortholog), the noise is high and we would, therefore, not recommend using this annotation level for any downstream analyses. The high noise at this annotation level compared to others is likely related to the inclusion of taxonomical information. Gene id annotation is composed of a tax id at the species level and the gene id from its original genome, making it challenging to accurately retrieve annotations from non-assembled reads.

Although not shown in this work, EggVio also integrates kaiju taxonomical annotation of the contigs, which is more accurate since it is carried out on longer reads, thus making the resolution much higher (as for functional annotation) and enables taxonomic analysis of the dominant strains of the population.

## 6 Conclusion

We showed that EggVio is a flexible pipeline for processing shallow depth sequencing datasets. In addition to assembling reads into contigs, the pipeline can rescue unrecruited reads while adding a predicted amount of noise (FDR = 5%) to the annotation. EggVio improved the correlation between metagenomic and environmental chemistry data. In addition, it removed the artifacts observed in the NMDS when calculated using the assembly alone and showed a more reliable community composition and allow a more robust sample comparison in the actual dataset. Nonetheless, further validation, using a mock community for example or benchmarking it against other possible tools such as miFaser is required to validate the full performance of the pipeline.

